# MeRIP-seq and RNA-seq data of mycelium and spore of *Ascosphaera apis*, a widespread bee fungal pathogen

**DOI:** 10.1101/2020.03.16.993840

**Authors:** Yu Du, Haibin Jiang, Zhiwei Zhu, Jie Wang, Huazhi Chen, Cuiling Xiong, Yanzhen Zheng, Dafu Chen, Rui Guo

## Abstract

*Ascosphaera apis* is widespread fungal pathogen of honeybee larvae, causing chalkbrood, a chronic disease that weakens bee health and colony productivity. In this article, mecylia and spores of *A. apis* were respectively purified followed by RNA isolation, cDNA library construction, MeRIP-seq and RNA-seq. A total of 62,551,172, 41,773,158, 49,535,092 and 61,569,610 raw reads were produced from Aam_IP, Aas_IP, Aam_input and Aas_input groups, respectively. After quality control, 58,484,368, 37,381,432, 44,655,434 and 58,739,742 clean reads were obtained. Furthermore, 47,706,205, 31,356,690, 35,259,810 and 44,319,061 clean reads were mapped to the reference genome of *A. apis*, including 39,337,036, 26,731,957, 31,987,396 and 40,017,855 unique mapped reads, and 8,369,169, 4,624,733, 3,272,414 and 4,301,206 multiple mapped reads. Among them, 96.31%, 96.51%, 96.82% and 97.11% of clean reads were mapped to exons; 2.09%, 2.31%,1.83% and 1.81% to introns; 1.60%, 1.18%, 1.35% and 1.08% to intergenic regions.

**Value of the data:**

- The data can be used to investigate the relationship between the m^6^A modification extent and the transcript level in the *A. apis* transcriptome.
- This dataset contributes to transcriptome-wide characterization of the m^6^A distributing patterns in mRNAs and non-coding RNAs in *A. apis* mycelium and spore.
- Current data benefits new functions of m^6^A modification in the transcripts extensively modified by m^6^A in *A. apis* mycelium and spore.
- Our data could be used to characterize differential patterns of the m^6^A methylation between mycelium and spore of *A. apis.*

## 1. Data Description

The shared data in this article were obtained from MeRIP-seq and RNA-seq of *A. apis* mycelia and spores. In total, 62,551,172 and 41,773,158 raw reads were respectively generated from Aam_IP group and Aas_IP group, while 49,535,092 and 61,569,610 raw reads were respectively produced from Aam_input group and Aas_input group (**Table 1**). After quality control, 90.06% and 84.33% clean reads in Aam_IP and Aas_IP were gained, respectively, with Q30 of 92.33% and 91.25% and GC content of 48.74% and 50.51% (Table 1); 68.45% and 77.07% clean reads in Aam_input and Aas_input were gained, respectively, with Q30 of 88.10% and 90.02% and GC content of 49.93% and 53.75% (**Table 1**). Further, 47,706,205 (81.64%) and 31,356,690 (84.05%) of clean reads in Aam_IP and Aas_IP were mapped to the reference genome (**Table 2**); among these 39,337,036 (67.32%) and 26,731,957 (71.65%) were unique mapped clean reads and 8,369,169 (14.32%) and 4,624,733 (12.40%) multiple mapped clean reads (**Table 2**). In Aam_input and Aas_input, 35,259,810 (79.95%) and 44,319,061 (76.55%) clean reads were mapped to the reference genome, including 31,987,396 (72.53%) and 40,017,855 (69.12%) unique mapped clean reads, and 3,272,414 (7.42%) and 4,301,206 (7.43%) multiple mapped clean reads (**Table 2**). Additionally, as **Figure 1** shown, 96.31% and 96.51% of clean reads in Aam_IP and Aas_IP were mapped to exons in *A. apis* genome, while 2.09% and 2.31% to introns and 1.60% and 1.18% to intergenic regions. In Aam_input and Aas_input, 96.82% and 97.11% of clean reads were mapped to exons, while 1.83% and 1.81% to introns and 1.35% and 1.08% to intergenic regions (**Figure 1**).

**Table 1.** Overview of raw reads and clean reads yielded from MeRIP-seq and RNA-seq.

| Sample | Raw reads | Clean reads | Ratio of clean reads | Q30 | GC content |
| --- | --- | --- | --- | --- | --- |
| Aam_IP | 62,551,172 | 58,484,368 | 90.06% | 92.33% | 48.74% |
| Aas_IP | 41,773,158 | 37,381,432 | 84.33% | 91.25% | 50.51% |
| Aam_input | 49,535,092 | 44,655,434 | 68.45% | 88.10% | 49.93% |
| Aas_input | 61,569,610 | 58,739,742 | 77.07% | 90.02% | 53.75% |

**Table 2.**
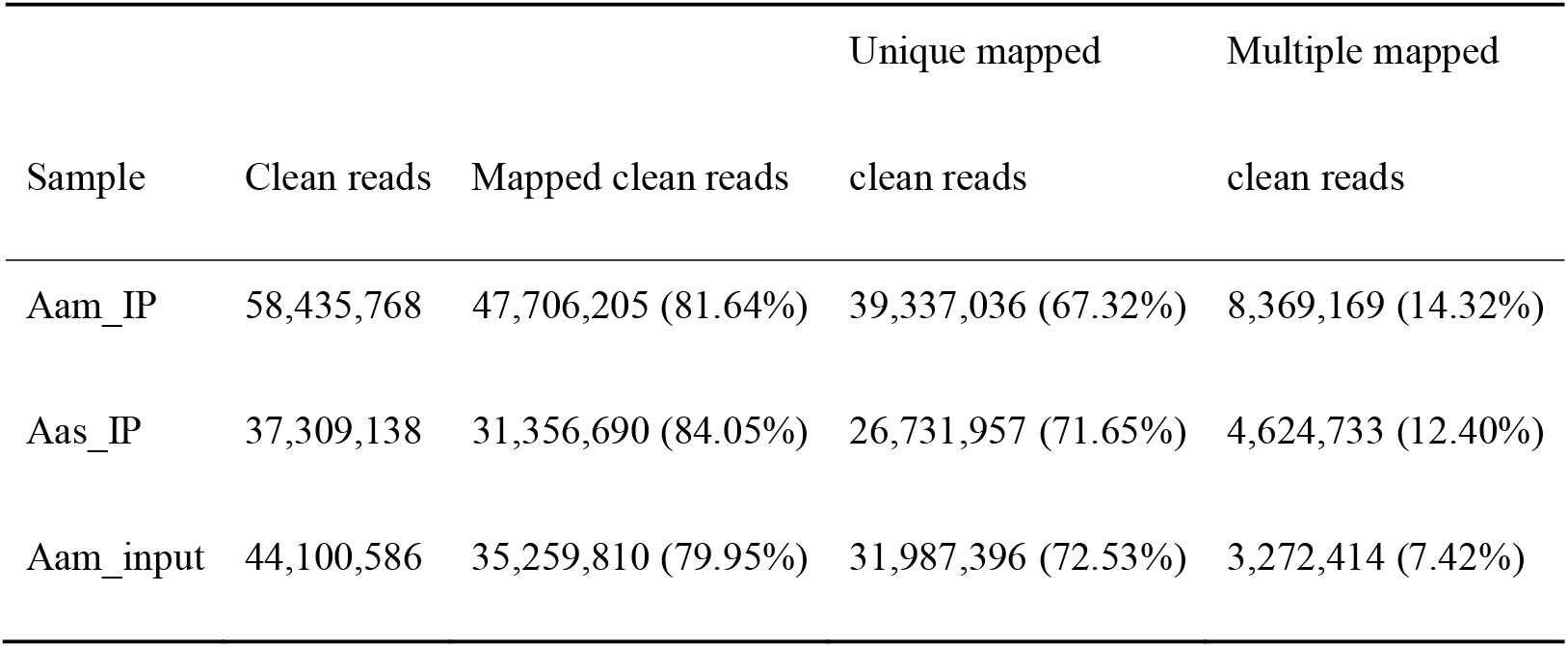
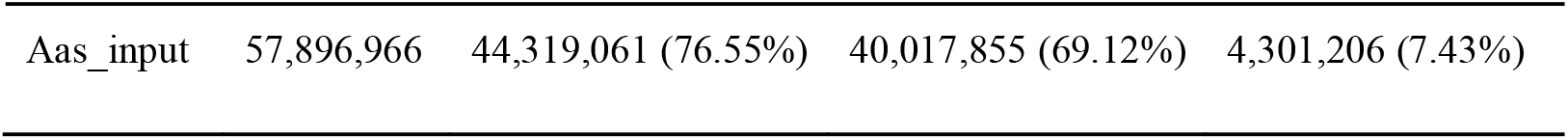
Summary of mapping clean reads to reference genome of *A. apis*.

**Figure 1.**
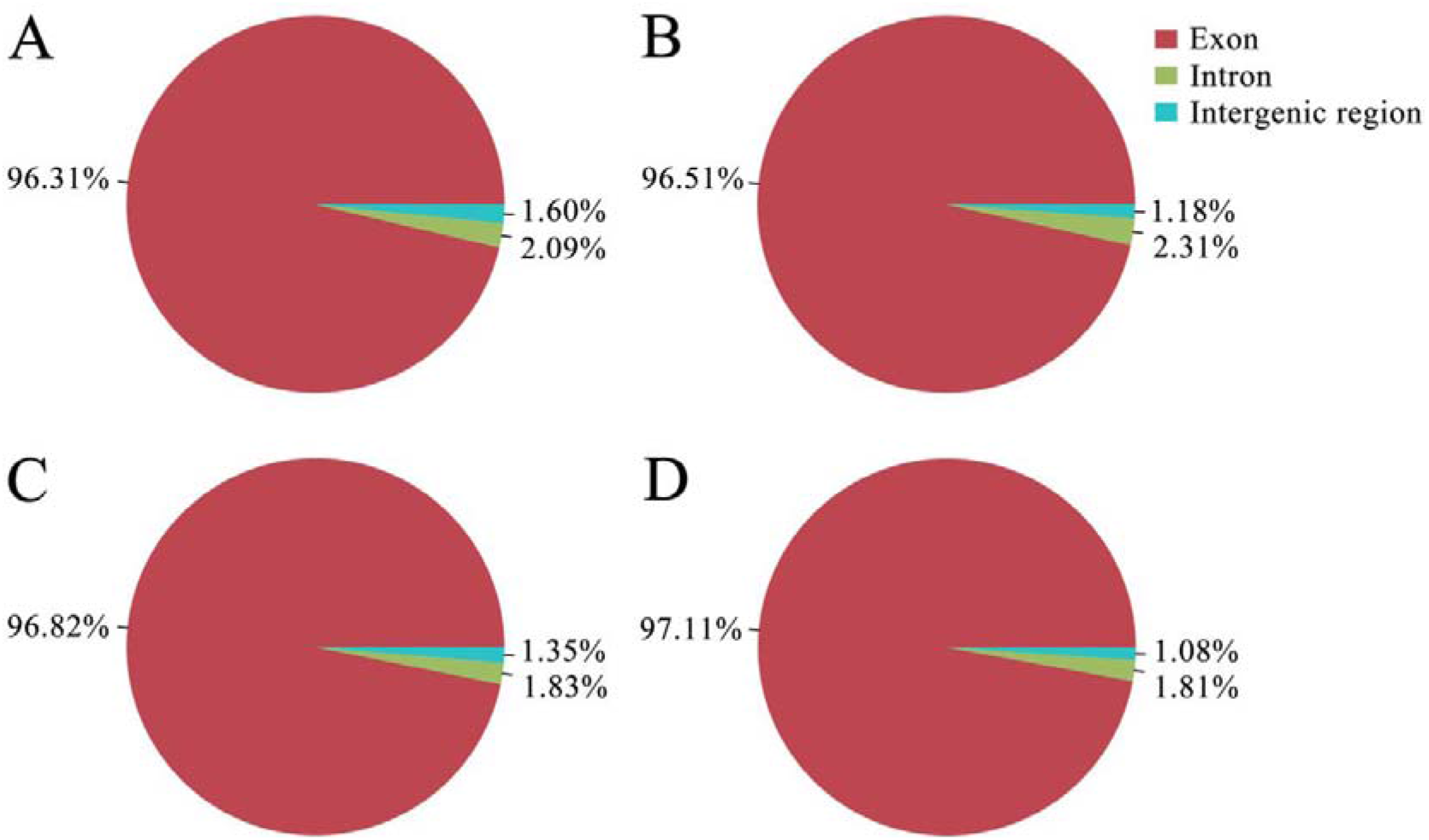
Genetic regions mapped by clean reads derived from *A. apis* mycelium and spore. A-D: mapping information of clean reads from Aam_IP, Aas_IP, Aam_input and Aas_input

## 2. Experimental Design, Materials, and Methods

### 2.1. Sample collection

*A. apis* was previously isolated from a fresh chalkbrood mummy of *Apis mellifera ligustica* larva [1] and kept at the Honeybee Protection Laboratory of the College of Animal Sciences (College of Bee Science) in Fujian Agriculture and Forestry University. *A. apis* mycelia and spores were respectively purified based on the previously developed method by Jensen et al. [2] with some modifications [3]. After purification, mycelium sample and spores sample were immediately frozen in liquid nitrogen and stored at −80 °C until MeRIP-seq and RNA-seq.

### 2.2. RNA isolation, library preparation, MeRIP-seq and RNA-seq

Firstly, total RNA was extracted using Trizol reagent (Invitrogen, USA) following the manufacturer’s procedure. RNA purity and quantification were evaluated using the NanoDrop 2000 spectrophotometer (Thermo Scientific, USA). RNA intergrity was assessed using the Agilent 2100 Bioanalyzer (Agilent Technologies, USA). Approximately more than 200 μg of total RNA was subjected to isolation of Poly(A) mRNA with poly-T oligo attached magnetic beads (Invitrogen, USA). Secondly, the poly(A) mRNA fractions were fragmented after purification, the fragments derived from mycelium and spores were divided into two parts: one was captured with m^6^A antibody to enrich mRNAs with m^6^A modification and the other was used as input for transcriptomic library construction. Thirdly, the RNA fragments used in MeRIP-seq were subjected to incubation at 4◻ for 2 h with m^6^A-specific antibody (No.202003, Synaptic Systems, Germany) in IP buffer (50◻mM Tris-HCl, 750◻mM NaCl and 0.5% Igepal CA-630) supplemented with BSA (0.02M DTT, 0.150M NaCl, 0.05M Tris–HCl (pH7.5), 0.001M EDTA, 0.10% SDS). Fourthly, the mixture was then incubated with protein-A beads and eluted with elution buffer (1×IP buffer and 6.7mM m^6^A). Eluted RNA was precipitated by 75% ethanol. Eluted m^6^A-containing fragments (IP) and untreated input control fragments are converted to final cDNA library in accordance with a strand-specific library preparation by dUTP method. Finally, the four libraries (Aam_IP, Aas_IP, Aam_input and Aas_input) were sequenced on an Illumina Novaseq™ 6000 platform (Illumina, USA) by OE Biotechnology (Shanghai, China), and 150 bp paired-end reads were generated.

### 2.3. Data processing

Firstly, Cutadapt software [4] and perl scripts in house were used to remove the reads that contain adaptor contamination, low quality bases and undetermined bases. Subsequently, quality of clean reads was verified using fastp. Next, HISAT2 [5] was used to map clean reads to the reference genome of *A. apis* (assembly AAP 1.0) with default parameters.

## Acknowledgments

This research was supported by the Earmarked Fund for China Agriculture Research System (No. CARS-44-KXJ7), the Science and Technology Planning Project of Fujian Province (No. 370 2018J05042), the Teaching and Scientific Research Fund of Education Department of Fujian Province (No. JAT170158), the Outstanding Scientific Research Manpower Fund of Fujian Agriculture and Forestry University (No. xjq201814), and the Scientific and Technical Innovation Fund of Fujian Agriculture and Forestry University (No. CXZX2017342, No. CXZX2017343).

